# RanDeL-seq: A high-throughput method to map viral cis- and trans-acting elements

**DOI:** 10.1101/2020.07.01.183574

**Authors:** Timothy Notton, Joshua J. Glazier, Victoria R. Saykally, Cassandra E. Thompson, Leor S. Weinberger

## Abstract

It has long been known that noncoding genomic regions can be obligate *cis* elements acted upon *in trans* by gene products. In viruses, *cis* elements regulate gene expression, encapsidation, and other maturation processes but mapping these elements relies on targeted iterative deletion or laborious prospecting for rare, spontaneously occurring mutants. Here, we introduce a method to comprehensively map viral *cis* and *trans* elements at single-nucleotide resolution by high-throughput random deletion. Variable-size deletions are randomly generated by transposon integration, excision, and exonuclease chewback, and then barcoded for tracking via sequencing (i.e., <u>**Ran**</u>dom-<u>**De**</u>letion <u>**L**</u>ibrary <u>**seq**</u>uencing, RanDeL-seq). Using RanDeL-seq, we generated and screened >23,000 HIV-1 variants to generate a single-base resolution map of HIV-1’s *cis* and *trans* elements. The resulting landscape recapitulated HIV-1’s known *cis*-acting elements (i.e., LTR, Ψ, and RRE) and surprisingly indicated that HIV-1’s central DNA flap (i.e., central polypurine tract, cPPT to central termination sequence, CTS) is as critical as the LTR, Ψ, and RRE for long-term passage. Strikingly, RanDeL-seq identified a previously unreported ∼300bp region downstream of RRE extending to splice acceptor 7 that is equally critical for sustained viral passage. RanDeL-seq was also used to construct and screen a library of >90,000 variants of Zika virus (ZIKV). Unexpectedly, RanDeL-seq indicated that ZIKV’s *cis*-acting regions are larger than the UTR termini, encompassing a large fraction of the non-structural genes. Collectively, RanDeL-seq provides a versatile framework for generating viral deletion mutants enabling discovery of replication mechanisms and development of novel antiviral therapeutics, particularly for emerging viral infections.

**Importance:** Recent studies have renewed interest in developing novel antiviral therapeutics and vaccines based on defective interfering particles (DIPs)—a subset of viral deletion mutant that conditionally replicate. Identifying and engineering DIPs requires that viral *cis*- and *trans*-acting elements be accurately mapped. Here we introduce a high-throughput method (Random Deletion Library sequencing, RanDeL-seq) to comprehensively map *cis-* and *trans*-acting elements within a viral genome. RanDeL-seq identified essential cis elements in HIV, including the obligate nature of the once-controversial viral central poly-purine tract (cPPT) and identified a new *cis* region proximal to the Rev responsive element (RRE). RanDeL-seq also identified regions of Zika virus required for replication and packaging. RanDeL-seq is a versatile and comprehensive technique to rapidly map cis and trans regions of a genome.

## Introduction

A generalized feature of genome structure is the presence and interplay of *cis*-acting and *trans*-acting elements (1). In viruses, trans-acting elements (TAEs) comprise viral gene-expression products such as proteins and RNAs that drive molecular processes involved in viral replication, maturation, and release (2). Viral cis-acting elements (CAEs) are sequences within the viral genome that are acted upon by TAEs, or that interact with other regions of the viral genome, to enable TAE-mediated genome replication, encapsidation, and other processes essential to viral maturation (3, 4). Across viral species, CAEs are conserved at the 5’ and 3’ ends, forming secondary structures such as stem loops and higher order structures that aid genomic stability or increase interaction with TAEs (5). Function can be often inferred from location, with 5’ CAEs correlating to replication and initiation of translation, and 3’ CAEs to nuclear export, RNA processing and RNA stability (6). CAEs can also be found within gene-coding regions and function in ribosomal frameshifting, RNA replication, and specifying the RNA for encapsidation (5).

Mapping and characterization of viral CAEs has elucidated critical molecular mechanisms in the lifecycles of a number of viruses (3, 7). For example, packaging signals, frameshifting signals, and internal ribosome entry sites (IRES) are critical CAEs and represent putative inhibition targets (8). Despite the challenges associated with disruption of structural elements, the high conservation rate of these sequences makes them attractive antiviral targets (4, 9).

One area where mapping of viral CAE and TAEs is clearly important is in rational design of live-attenuated vaccines (LAVs) (10, 11); LAV-candidates lacking CAEs have reduced replicative fitness. Thus, CAE retention may be required for efficient replication and immunogenicity of the LAV candidate. Alternatively, it is possible that deletion of CAEs could enable calibration of viral replication for attenuation. Knowledge of conserved features is also important for viruses subject to high recombination or mutation rates (12), and a rapidly implementable, attenuation platform would clearly be beneficial (13). Additionally, knowledge of conserved viral regions aids the development of complementary attenuation strategies, such as microRNAs (14).

Mapping viral CAEs and TAEs may also aid development of novel classes of antivirals that act via genetic interference (15) and are proposed to have high barriers to the evolution of viral resistance. One class of proposed antivirals are Therapeutic Interfering Particles (TIPs), engineered molecular parasites of viruses based upon Defective Interfering Particles (DIPs). DIPs are sub-genomic deletion variants of viruses that do not self-replicate but conditionally mobilize in presence of wild-type virus and can interfere with wild-type replication (16-23). TIPs are enhanced DIPs proposed to retain all CAEs and interfere with wild-type replication by stoichiometric competition for TAEs, such as packaging proteins, within the infected cell. Enhanced replication of DIP/TIPs in turn reduces the wild-type viral load. Current candidates (24-26) are generated by traditional methods of high-multiplicity of infection (MOI) serial passage or UV-inactivation. A high-throughput, rational genetics approach to development DIPs and TIPs would aid screening and identification of safe and effective candidates.

Despite the benefits of mapping viral CAE and TAEs, methods to do so, especially for CAEs, tend to be laborious and/or highly technical, and traditionally focused on protein-coding sequences, rather than on regulatory sequences (27, 28). Highly technical methods include multicolor long-term single-cell imaging (29), CRISPR/Cas9 deletion tiling (27, 30), chemical probing approaches (31), targeted RNA mutagenesis and functional binding assays (32, 33), and bioinformatics (6, 34). Most methods, however, still rely on viral defective interfering (DI) RNAs, deducing critical genomic regions by serial passage. CAEs are in turn mapped by analyzing deletion variant sequences that persist or produce infectious virions. DI RNA studies reveal critical genomic regions that can be investigated further with reverse genetics systems such as site-directed mutagenesis (35-39) or iterative deletion vectors (40-44).

These approaches are limited by the ability to examine one element at a time, iterating deletions around one factor, or deleting portions of the viral genome. Not all viruses have naturally occurring DI RNAs, and generating them by serial passage is straightforward, but laborious. Deletion mutants arise at low frequency and remain rare unless a deletion confers increased fitness relative to the wild-type virus. A number of methods to generate defined mutants and random deletions at an appreciable frequency using reverse genetic systems exist, such as creating short random deletions with endonucleases (45) or using synthetic DNA and site-specific recombinases (46) for larger deletions. Other methods insert transposon cassettes into viral genomes to disrupt CAE and TAEs by separating protein domains or introducing missense and nonsense mutations (47-50). These methods do not generate deletion mutants at scale, and all have certain drawbacks, whether it be non-random mutation/deletion, viral insertion scarring, reliance on previously characterized DIPs, inability to generate and track full-length viral mutants, or the price, labor, and versatility of the method.

In this study, we present a versatile framework for generating random deletion libraries of viral species in high throughput and mapping viral CAE and TAEs without laborious and iterative deletion. Through *in* vitro transposition, dual exonuclease chewback, and barcode ligation, RanDeL-seq (<u>**Ran**</u>dom <u>**De**</u>letion <u>**L**</u>ibrary <u>**seq**</u>uencing) generates diverse randomized libraries of barcoded viral deletion variants (>10^5^ unique mutants) at modest expense in fewer than 5 days. As proof of concept, we demonstrate the construction and screening of tagged libraries of >23,000 deletion mutants of HIV-1 and >90,000 deletion mutants of Zika virus (ZIKV). Repeated *in vitro* passage and deep sequencing of the pooled viral mutants comprehensively mapped HIV-1 and ZIKV at single-base resolution, identifying established viral CAEs and revealing the importance of other viral regions for sustained viral replication in cells, such as the cPPT and splice acceptor 7 (SA-7) in HIV-1 and non-structural proteins in ZIKV.

### A Method to Generate a Random Deletion Library (RanDeL) – HIV-1 case study

To map viral cis- and trans-acting elements, we developed RanDeL-seq, a technique to efficiently generate and screen <u>**Ran**</u>dom <u>**De**</u>letion <u>**L**</u>ibraries of a viral species in high-throughput. The method (Fig 1A) involves deletion via *in vitro* transposition, transposon excision, dual exonuclease chewback, and re-ligation with molecular barcodes able to be mapped by deep sequencing. Viral mutants could be followed over time by their unique barcodes at a resolution not attainable by standard sequencing of pooled viral nucleic acids (51).

**Figure 1:**
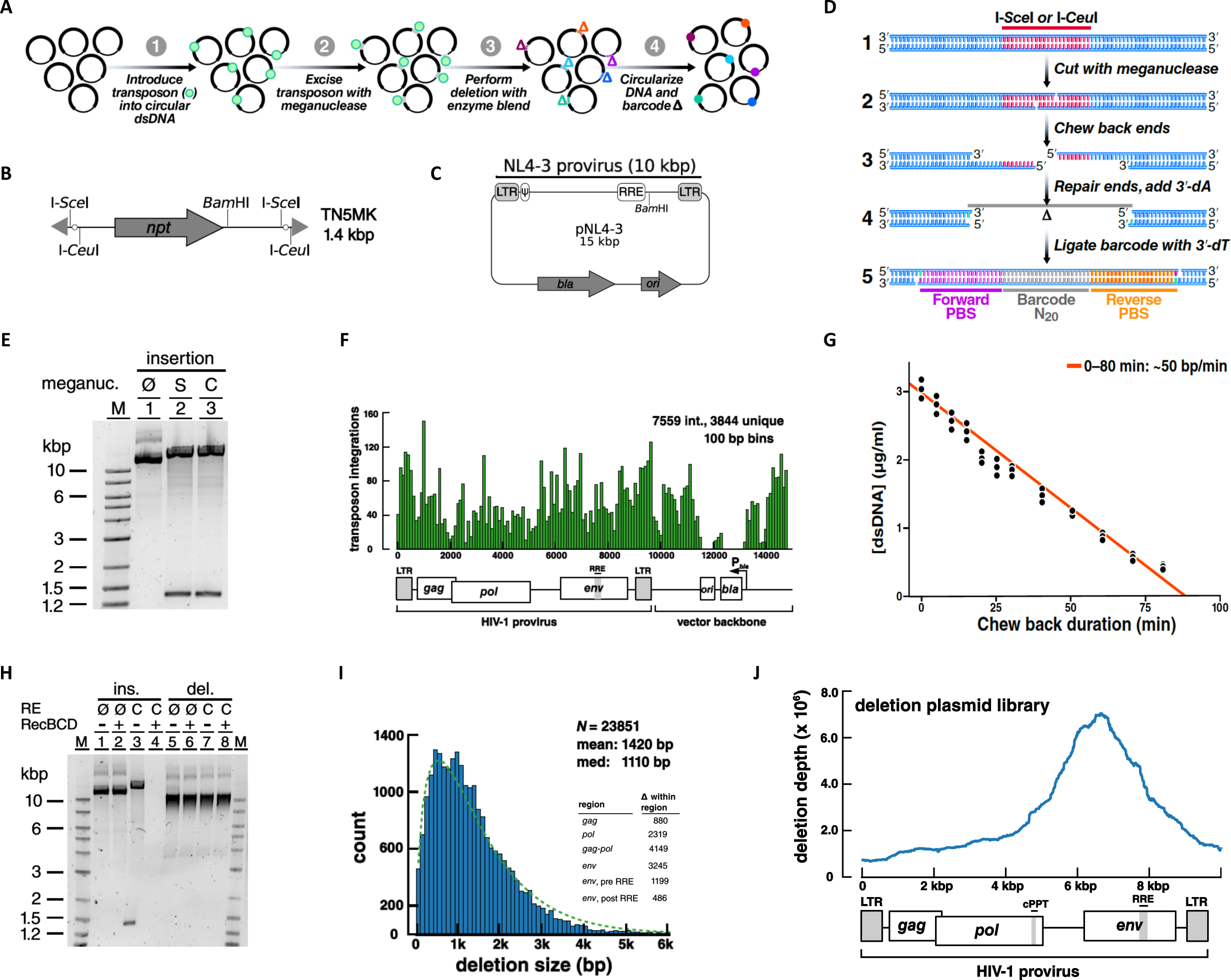
Method to generate a Random Deletion Library in HIV-1. A. Overview schematic of method to create a barcoded random deletion library: (1) Transposon cassettes harboring unique restriction sites are inserted into plasmids via in vitro transposition. (2) Transposons are excised to linearize the insertion library with a meganuclease. (3) Deletions are performed by chewback from both DNA termini by simultaneous treatment with enzyme blend. Mean deletion size is modulated by adjusting duration of chewback. (4) The chewed termini are end-repaired, dA-tailed, then joined by ligation to a T-tailed 60bp unique barcode cassette. B. Schematic of the “TN5MK” synthetic meganuclease transposon cassette used in library construction: TN5MK is composed of an antibiotic resistance gene, neomycin phosphotransferase I (npt), flanked by meganuclease restriction sites for I-SceI and I-CeuI and Tn5 mosaic ends (gray triangles) at the termini. The transposon cassette also contains a unique internal BamHI recognition site. C. The HIV-1 molecular clone pNL4-3, is a 14825 bp plasmid harboring the 9709 bp NL4-3 provirus (HIV-1 subtype B). NL4-3 is a chimera of two viruses (NY5 and LAV). D.Library insertion, excision, barcoding details: Circular DNA (1) is linearized by digestion with a meganuclease (I-SceI or I-CeuI), which cleaves at recognition sites encoded on the inserted transposon. This creates linear DNA with 4 base 3’ overhangs (2). Deletions are created by bidirectional chewback. Treatment with two exonucleases (T4 and RecJf) creates a population of truncated deletion mutants with ragged ends (3). Ragged DNA ends are blunted and then prepared for barcode cassette ligation by 5’ dephosphorylation and addition of a single 3’-dA (4). Deletion mutants are re-ligated in presence of a barcode cassette with single 3’-dT overhangs and 5’ phosphoryl groups to create barcoded circular DNAs with 2 nicks separated by 60 bp (5). E. Insertion libraries following I-SceI (S) or I-CeuI (C) digestion. Digestion of pNL43 insertion library shows excisions of the TN5MK transposon (1.4kb) and upward shift of the supercoiled library vs. the undigested library. Lanes: (M) 2 log DNA ladder, (1) undigested insertion library, (2) I-SceI digested insertion library, (3) I-CeuI digested insertion library. F. Location of TN5MK insertions for a subset of 7559 transposon integrations (3844 were unique). G.Determination of enzymatic chewback rate for deletion size: The chewback rate was determined by treating a 4 kb fragment of linear dsDNA with RecJf and T4 exonucleases in the presence of SSB and no dNTPs for increasing amounts of time, then halting enzymatic activity. Reactions were performed in triplicate. DNA concentrations were established by quantifying fluorescence of PicoGreen in a plate reader in comparison to a dsDNA standard of known concentration. H.Validation of Deletion Library: The pNL4-3 insertion library and pNL4-3 deletion library were either not digested (Ø) or cut with I-CeuI (C) and then subjected to binary treatment with RecBCD, which digests linear DNA to completion. Lanes 1-4 are the pNL43 insertion library and Lanes 5-8 are the pNL43 deletion library. I.pNL4-31 is composed of 23,851 tagged mutants with a range of deletion sizes. The right-skewed (i.e. right-tailed) histogram of deletion sizes in pNL4-31, with bins of 100 bp (shown in blue) is well-fit by a Gamma distribution (green, broken-line). Inset: Number of deletions detected within each region of the HIV genome. J.Deletion Depth Profile over the full HIV-1 genome. Calculation of the deletion depth profile of the pNL4-31 genome indicates that each base is covered by hundreds to thousands of deletion mutants. Two regions where deletions are not tolerated in the plasmid backbone are ori, the origin of replication and bla, -lactamase, the resistance marker

To start, we designed a synthetic transposon cassette, TN5MK, (Fig. 1B) compatible with the well-characterized hyperactive Tn5 transposase (52-54). Transposons contained an antibiotic-resistance marker to select for plasmids harboring a successful transposon insertion. Transposon integration into the target plasmid introduces unique restriction sites for uncommon meganucleases, I-SceI and I-CeuI, with long recognition sites (Fig S1A). The length of the recognition site confers specificity and is advantageous for use without modification in many systems.

The conventional HIV-1 molecular clone pNL4-3 (Fig. 1C), was the substrate for this library construction. The system allows control over the size of deletions and can tag each member of the diverse deletion library with a molecular barcode to facilitate deep sequencing analysis. The molecular biology details of RanDeL-seq are in Figure 1D. Each step was validated after completion, with a comprehensive check on the finished libraries.

First, we performed *in vitro* transposition to randomly insert TN5MK into pNL4-3 at a ratio of one transposon per viral plasmid. The insertion libraries were treated with both encoded restriction enzymes, generating the expected ∼ 1.4 kb excised transposons in addition to linearized pNL4-3 plasmid backbone (Fig. 1E). Deep sequencing of the pNL4-3 insertion library enabled mapping of insertion sites across the genome (Fig. 1F), showing TN5MK integrated broadly throughout the pNL4-3 plasmid, with a high frequency and density of at least one transposon integrated every 100 bp. In the plasmid backbone, two integration gaps emerged: one in origin of replication, ori, and the other in the resistance marker, as both are required for propagation of the plasmid in *E. coli*.

After creating a polyclonal population of transposon-inserted circular target DNAs, insertions were excised by meganuclease treatment. DNA chewback with a trio of proteins (T4 DNA polymerase, RecJF, and SSB) efficiently created truncations in a common buffer system (Fig S1B). Chewback rate was determined by a dsDNA fluorometric method, using a 4 kb template DNA. As the ends were progressively shortened by chewback, the fluorescence signal of a dsDNA-specific dye (PicoGreen) decreased proportionally. The measured double-end chewback rate, as determined by linear regression, was approximately 50–60 bp/min (Fig 1G). Sub-libraries of mutants with diverse deletion sizes were then created by varying the enzymatic incubation time. Finally, linearized sub-libraries were pooled, end-repaired, dA-tailed, dephosphorylated, and recircularized by ligation to a 3’ dT-tailed 60 bp barcode cassette. The barcode cassette was designed to have a 20-bp random barcode flanked by 20-bp primer-binding sites, taken from Tobacco Mosaic Virus to limit sequence complementarity with human viruses (Fig S2). Each successful ligation resulted in a deletion mutant tagged with a unique barcode cassette.

We validated the final library via several different methods (Fig. 1H). First, to test if the transposon insertion was fully excised, libraries were restriction enzyme digested by I-CeuI. The completed library (Lane 7) was insensitive, compared to a digested insertion library (Lane 3), confirming TN5MK excision and removal in chewback. Second, an untreated deletion library had a range of sub-genomic sizes (Lane 5) in comparison to an untreated insertion library (Lane 1), confirming that chewback created deletions of various sizes. Lastly, to test successful recircularization with the barcode cassette, the digested insertion and deletion libraries were treated with RecBCD, an enzyme that degrades linear dsDNA. We hypothesized that treated insertion libraries and deletion libraries that maintained I-CeuI cut sites or were not properly ligated to barcode cassettes will be completely degraded. Post-treatment the deletion library was unchanged, as the digested plasmids are uncut and circular from ligation to the barcode cassette (Lanes 6 & 8). On the other hand, treatment of insertion libraries degraded all plasmid (Lane 4).

### Framework for efficiently sequencing barcoded HIV-1 RanDeL

Post validation of the tagged RanDeL, we developed a framework for genotyping barcoded mutants in order to track each unique deletion mutant in culture and calculate a viral deletion depth profile. RanDeL-seq relies on the initial whole genome sequencing of the deletion library to construct a look up table that links each unique barcode sequence to a specific deletion locus. This initial genotyping step allows for efficient sequencing downstream, as only barcode cassettes need to be sequenced downstream to identify which deletion mutants persist in culture.

The plasmid library was fragmented, deep-sequenced (2 × 125b reads on HiSeq4000) with Illumina™ paired-end sequencing and analyzed with custom python software (rdl-seq). Reads were filtered for the small percentage (2.9%) that contained the full barcode cassette (Table S1). Repeated barcode sequences were grouped together to determine the consensus bases 5’ and 3’ of that specific barcode cassette (i.e., barcode flanking sequences). Flanking sequences were aligned to the viral reference genome, generating a lookup table of barcodes (B = b_1_, b_2_, b_3_ … b_n_) matched to deletion loci (D = d_1_, d_2_, d_3_… d _n_). After the initial genotyping of the plasmid random deletion library, deletion variants can be identified by amplifying barcodes cassettes with primers annealing to the common primer-binding sites.

This sequencing framework determined there were 23,851 unique mutants with a range of deletion sizes (Fig 1I). The library subset had a median deletion size of ∼ 1.1 kb, a minimum deletion of 30 bp and a maximum deletion size of >6 kb. The skewing of the library (i.e. long tail to high kb), may be due to mechanical shearing of DNA during some of the cleanup steps.

Next, the deletion-depth profile (location and abundance of deletions) of the pNL4-3 deletion library was calculated (Fig 1J). The plasmid library exhibits deletions across the HIV genome, with a peak in the *env* gene, and a region of zero deletion depth was observed at the plasmid origin of replication (ori) and antibiotic resist marker (bla). Biases in the deletion depth at this stage corresponds to differences in bacterial growth rate; faster growing bacteria lead to overrepresentation of their harbored plasmid. The signal peptide of *env* and sequences at the N-terminus are known to be toxic to bacteria, therefore bacteria harboring *env* deletions likely have a growth advantage, and bacteria harboring ori/bla deletion plasmids are unable to grow in the antibiotic.

### Serial-passage screening of HIV-1 RanDeL to map viral CAE and TAEs

To functionally characterize the deletion library, we designed a high multiplicity of infection (MOI) passage scheme to select for and map viral CAEs by sequencing the barcodes that persisted through multiple passages together with replication-competent HIV-1. A high MOI ensured that on average, each cell became infected with more than one copy of the wild-type virus to supply trans factors. The diversity of the library was limited to fewer than the number of available cells to maintain strong selective pressure, avoid drift, and ensure that most of the library would be sampled multiple times during infection. In this scheme, the genomic regions that can tolerate deletion (as measured by enrichment of specific barcodes corresponding to that region) correspond to TAEs, while regions that are intolerant of deletion correspond to CAEs. Regions of the genome could also be neutral (i.e., extraneous or ‘junk’ regions) but given the extreme selection pressures that viral genomes face, such neutral regions (non cis, non trans) are expected to be small, especially for RNA viruses.

Wild-type virus and deletion library pools were packaged by co-transfection of 293T cells with equal masses of the pNL4-3 deletion library and pNL4-3 parental plasmid. Clarified supernatant (0.45 um filtered) was concentrated by ultracentrifugation and used to transduce MT-4 cells at high MOI in three parallel biological replicates (designated K, L, and M) for twelve passages (Fig 2A). In parallel, three flasks were infected with wild-type HIV-1 only as a negative control for deletion library barcodes. In this high MOI passage scheme (Fig 2B), cultures were infected with concentrated virus, then supplemented with naive MT-4s every 24 hours before being harvested 3 days (i.e. 3 passages) later. By supplying naive target cells, the scheme selected for two phenotypes: (a) replication-competent viruses and (b) replication-defective viruses that are efficiently trans-complemented by wild-type virus (i.e., mobilized). Flow cytometry of high MOI conditions showed an initial high percentage of infected cells, followed by an expected drop after the addition of naive MT4s, and then a return to high infected percentage before harvest. (Fig S3).

**Figure 2:**
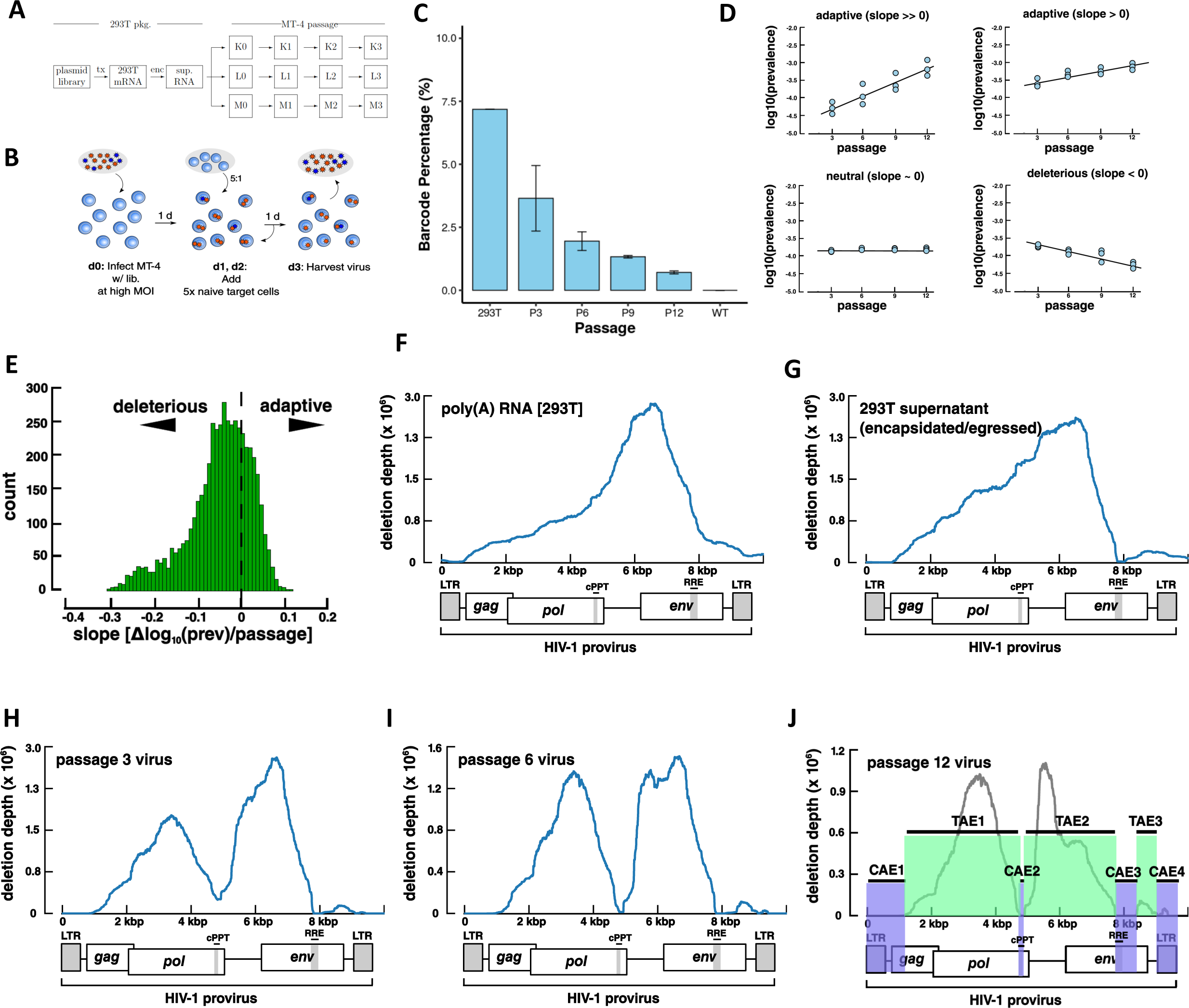
Genetic Screen of random deletion library to map viral cis- and trans- elements. A. Block design of high-MOI passage. Wild-type NL4-3 and deletion library plasmids co-transfected 293T cells. Virus-containing supernatant infected MT-4 cells in triplicate (K, L, M) at high MOI. Infections were passaged at the end of every week, for 4 weeks. At the same time, flasks with only NL4-3 wild-type virus were passaged identically (A, B, C not shown). B. Passage details of high MOI screen. MT-4 cells (blue double discs) are infected at high MOI with a pool of virus (HIV-1) containing both wild-type (red stars) and deletion mutants (blue stars). At days 1 and 2, additional naïve MT-4 were added and the culture volume expanded. On day 3, cell-free supernatant was harvested, and virus purified by ultracentrifugation for transfer or analysis. One passage corresponds to 3 rounds of replication. C. Detection and quantification of barcode cassettes by RT-qPCR. Genomic percentage of barcoded mutants to total HIV genomes in transfection (293T), each stage of high MOI passage (P3-P12), and a wild-type HIV control (WT). RT-qPCR data was normalized to a MS2 RNA spike-in. Error bars are standard deviation from averaging each flask (K, L, M) per passage. D. Representative deletion variant trajectories during high MOI passage. The slope in prevalence versus passage number was determined by linear regression and classified deletions as adaptive, neutral or deleterious. Data points correspond to the triplicate flasks (K, L, M) at each passage. Prevalence is in reference to the total barcode cassette pool (tagged mutants). E. Distribution of fitness in deletion variants that are not extinct by passage 12. The dashed vertical line marks the neutral fitness boundary (slope of 0). 1390 of 4390 persistent mutants were adaptive. F. The deletion depth profile of poly(A) RNA from transfected 293T cells, representing mutants able to be transcribed. G. The deletion depth profile built from the virus-containing supernatant of transfected 293T representing mutants able to be transcribed, encapsulated, and egressed. H. Deletion depth profile from virus-containing supernatant after 3 passages. I. Deletion depth profile after 6 passages. J. A model of HIV-1 cis- and trans-acting elements after 12 passages. The HIV-1 genome is composed of 4 cis-acting elements, CAE1–CAE4, (highlighted in blue) and 3 trans-acting elements, TAE1–TAE3, (highlighted in green).

### Tracking RanDeL barcodes throughout serial-passage experiments

Viral RNA from cell-free supernatants was analyzed by RT-qPCR to detect barcode sequences and determine which deletion variants persisted passage to passage. Barcodes were detectable in all deletion library samples in the serial-passage, and in none of the control flasks. The ratio of barcodes to total HIV genomes slightly decreased over time from the initial co-transfection of deletion library samples (Fig. 2C). Expression of total HIV genomes was not significantly different between the library and control samples (Fig S4A), indicating no interference from the deletion library.

Using custom Illumina-prep for barcode sequencing (Fig S5), the prevalence of each deletion variant in the total population of barcoded mutants was tabulated, and the prevalence trajectory throughout the passage computed. Of the 23,851 mappable pNL4-3 deletion mutants, only 4390 (18%) were detectable in all three replicate flasks by passage 12—the remaining 19,461 (82%) barcodes were undetectable and presumably were extinct in at least one of the three replicates. Overall there was strong concordance in barcode prevalence between the three replicates (Fig S4B).

We computed trajectories for the 4390 barcoded deletion variants, calculated the change in prevalence versus passage number (i.e., slope) by linear regression, and classified variants by slope (Fig 2D). Linear regression analysis determined that 1390 (32%) of the 4390 persisting deletion variants increased in prevalence through every passage, indicating that variants harboring these deletions were transmitting better than the average member of the barcoded population (Fig 2E). The remaining 3000 mutants remained steady or decreased in prevalence passage to passage. As barcode levels were relatively constant to total HIV genomes, we hypothesize that these 1390 persisting variants are transmissible (R0>1) and can be efficiently complemented in *trans* (i.e., these deleted regions can be compensated for by gene products expressed from wild-type HIV-1) and can spread through the population as fast or faster than wild-type HIV-1.

### HIV-1 Deletion Landscapes identify CAE and TAEs

Using deep-sequencing counts of barcodes and referencing back to the barcode-to-genotype look-up table, deletion landscapes (a.k.a. deletion-depth profiles) were calculated for the HIV-1 genome at various timepoints in the screen. First, we sequenced barcodes in intracellular poly(A) RNA purified from the 293T cells (Fig. 2F) used to package the deletion library. The 5’ end of the HIV-1 genome (spanning the 5’ LTR through SL1–SL4) exhibited low deletion depth, while the rest of the genome showed little reduction in barcode coverage. This deletion landscape reflects known CAEs required for efficient HIV-1 transcription in 293T cells.

Next, barcodes were sequenced from the cell-free supernatant of 293T cells (Fig. 2G), representing deletion variants able to be transcribed, packaged into virions (encapsidated), and released from the cell (egressed). This supernatant deletion landscape differed from intracellular RNA deletion landscape in two key genomic regions: (i) the region of zero deletion depth beginning at the 5’ LTR and extending through the start codon of *gag*, which includes the HIV-1 packaging signal (*psi*, Ψ), and (ii) at the 3’ end of the genome, the stretch of zero deletion depth that maps to the Rev Responsive Element (RRE)—a region of secondary structure critical for nuclear export of incompletely spliced HIV-1 RNAs (55, 56). These data indicate that the LTR, Ψ, and RRE were the only elements critical for efficient transcription, encapsidation, and egress of HIV-1 from 293T cells; all other regions tolerated some amount of deletion.

Deletion landscapes were then calculated to profile the changes in the deletion library passage to passage in MT-4 cells. At passage 3 (Fig. 2H), the deletion landscape diverged notably from the 293T-intracellular and supernatant profiles in three key ways: (i) a valley of reduced deletion depth appeared, with a minimum centered above the cPPT/CTS, (ii) the region of zero-deletion depth at the 5’ end of the genome shifted, encompassing the 5’ LTR through the first three hundred bases of *gag*, and (iii) a widening and 3’ shift of the deletion-depth valley situated around the RRE occurred. At passage 6 (Fig. 2I), these features had become more pronounced, and each valley had flattened to a deletion depth near zero.

No significant landscape differences were found between passages 9 and 12, enabling construction of a consensus map (Fig. 2J). Three regions of the HIV-1 genome were tolerant to deletion and able to be complemented efficiently in *trans*. These deletion-tolerant regions were classified as TAEs and are: (i) a region centered at the deletion peak at the center of *pol* (TAE1), (ii) a region in HIV’s accessory gene tract (*vif–vpu*) (TAE2), and (iii) a region in the 3’ end of *env* (TAE3).

The final deletion-depth profile contained four regions of low or zero deletion depth, indicating that these genomic are required CAEs. CAE1 is the first 1114 nucleotides of the proviral genome, encompassing known CAEs the 5’ LTR, stem loops 1–4, and the first 325 bp of *gag*, which maps to the Gag MA (p17) and Ψ. CAE2 maps directly to the cPPT/CTS. The requirement for HIV cPPT in reverse-transcription and integration has been debated in the past, with the literature supporting (31, 57-61) and questioning (62-65) its role. Here, the data support a critical role for cPPT in sustained HIV-1 replication. CAE3 begins at the RRE and ends precisely at splice acceptor 7 (SA-7), which is used for several multiply spliced HIV-1 transcripts, including vpr, tat, rev, nef (66), and implicated in viral fitness (56). While the importance of the RRE and SA-7 were known (31, 67, 68), RanDeL-seq showed that the entire 300 bp region from the upstream RRE to the end of SA7 is required for sustained viral replication, and cannot be provided in *trans*. Finally, CAE4 spans the PPT, which is necessary for reverse transcription, and the 3’ LTR (18).

### Application of RanDeL-seq to identify Zika Virus CAEs

To determine if this approach has the potential to be more generally applicable across diverse viruses, we performed RanDeL-seq on Zika virus (ZIKV). ZIKV is a flavivirus with a (+)-stranded, ssRNA genome of approximately 11 kb that replicates predominantly in the cytoplasm of infected cells. Libraries were built using two cDNA molecular clones of the conventional 1947 Ugandan strain of ZIKV, MR-766 (69). The first clone, Pol(+) pMR766, encodes the wild-type virus (Fig. 3A), whereas the second clone, Pol(-) pMR766, encodes a defective mutant with a substitution in the active site of the essential RNA dependent RNA polymerase NS5. Consequently, pMR766(-) virus is not replication-competent, unless rescued by providing NS5 in trans. pMR766(+) and pMR766(-) insertion libraries were generated with TN5MK. Next, the transposon was excised, enzymatic chewback performed to generate deletions, and the cDNA re-circularized by ligation in the presence of a random barcode cassette. Both short (S) and long (L) duration chewbacks were performed for each insertion library to create small and large average deletion sizes, respectively. Overall, four ZIKV RanDeLs were generated: pMR766(+)S, pMR766(+)L, pMR766(–)S, and pMR766(–)L.

**Figure 3:**
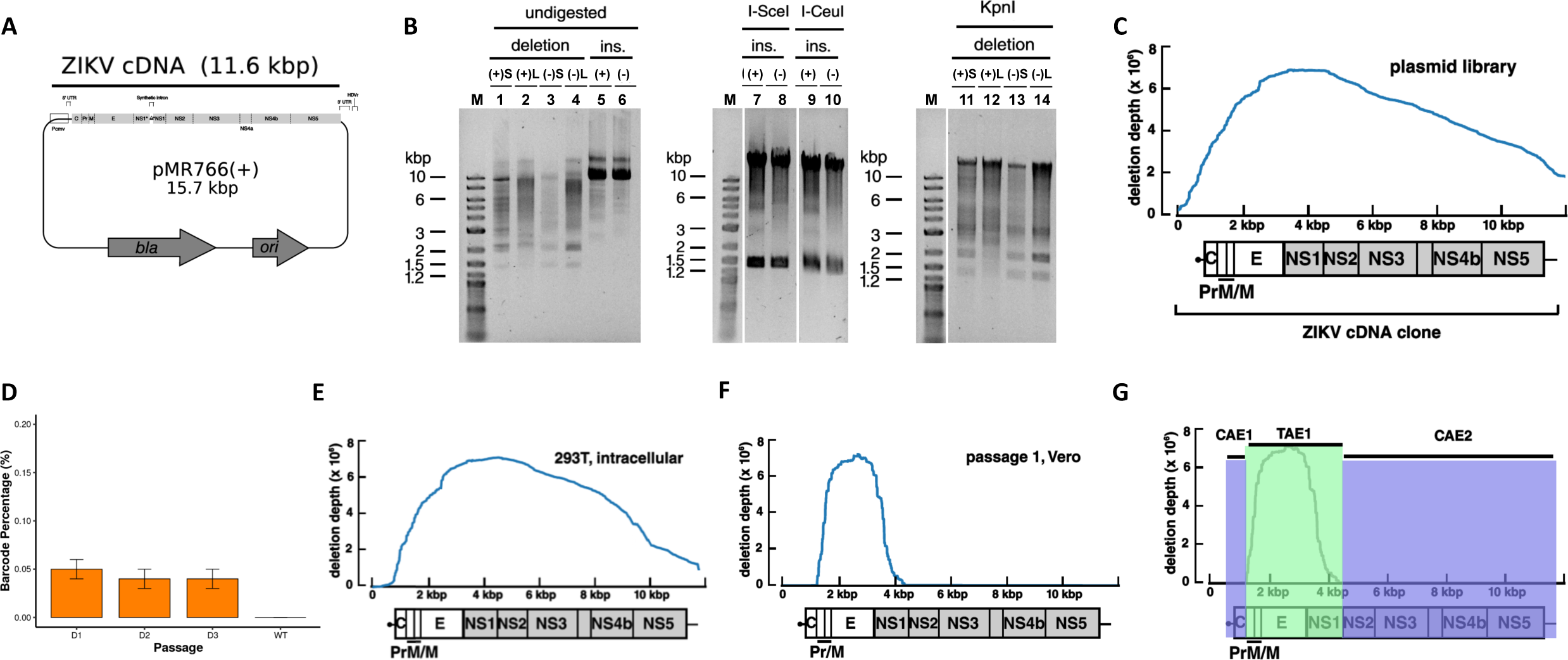
Application of RanDeL-seq to map Zika Virus (ZIKV) cis elements. A. pMR766(+), a Zika virus molecular clone. The MR766 Zika virus genome is encoded as a cDNA driven by the CMV IE2 promoter. At the 3’ end of the genome, a self-cleaving Hepatitis Delta ribozyme allows for creation of an authentic 3’ end post-transcription. An intron sequence is present within NS1 to allow maintenance in bacteria but is spliced out during transcription in host cells. B. Restriction enzyme characterization of completed ZIKV deletion libraries compared to insertion libraries (“ins.”). (+) and (-) designate the template ZIKV plasmid. “S” and “L” designate the chewback length for deletion libraries. Undigested, completed deletion libraries (Lanes 1-4) were run next to undigested insertion libraries (Lanes 5-6). Insertion libraries (Lanes 7-10) treated with I-SceI or I-CeuI to excise transposon (∼1.4 kb). Deletion libraries linearized by unique ZIKV cutter KpnI (Lanes 11-14). C. Deletion depth profile of the pMR766(+)L library. The ZIKV genome is well-represented in the pMR766(+)L library, with some bias. Each base of the ZIKV genome is covered by several hundred different deletion mutants. D. Detection and quantification of ZIKV barcode cassettes by RT-qPCR. Genomic percentage of barcoded mutants to total ZIKV genomes at each day in passage 1 of the high MOI screen, and a wild-type ZIKV control (WT). RT-qPCR data was normalized to a MS2 RNA spike-in. E. Deletion Depth Profile of intracellular RNA of 293T co-transfected with the wild-type ZIKV plasmid and the pooled deletion libraries. F. Deletion Depth Profile of pMR766(+)L after Passage 1. Only deletions in Pr–NS1 can be trans-complemented by wild-type ZIKV. H. Final map of ZIKV cis- and trans-acting elements after Passage 2. The two cis-acting regions are highlighted in blue and do not tolerate deletion (i.e., must be present for efficient transmission to occur). The trans-acting region is highlighted in green and can be complemented in trans (i.e., if deleted, transmission occurs by complementation from wild-type virus).

Each ZIKV RanDeL was validated per methods similar to those used for the HIV-1 library (Fig. 3B). Restriction-enzyme analysis with I-SceI and I-CeuI showed transposon excision in both sets (+ and -) of insertion libraries (lanes 7-10). Undigested, completed deletion libraries ran at 2–10kb (lanes 1–4) confirming various sized deletions as a result of chewback incubations. Successful plasmid re-ligation and re-circularization of each deletion library was analyzed by restriction analysis with KpnI, a restriction enzyme with a single unique site in both ZIKV wild-type plasmids. Consistent with successful plasmid re-circularization, KpnI digestion generated single bands (lanes 11-14)—i.e., linearized molecules arising from a cut of a circular plasmid as opposed to two molecules arising from cutting of a non-circularized, linear DNA molecule. These KpnI-digested single bands migrated at sizes larger than the undigested supercoiled libraries KpnI (lanes 1-4).

Whole-plasmid sequencing of the four ZIKV RanDeLs determined the deletion diversity to be between 1,000 and 50,000 mappable deletions per library (Table S1). Short-chewback libraries had less diversity than long-chewback libraries, likely due to some short-chewback reactions failing to chew past the transposon cassette, rendering it impossible to determine the mutation location. The deletion-size distribution of the ZIKV RanDeLs differed from the HIV-1 RanDeLs (Fig. S6) in that ZIKV RanDeL distributions were clearly bimodal, with peaks at small and large deletion sizes. Increasing the length of the chewback shifted the lower peak, but not the upper peak, possibly indicating that clones that lost the ZIKV cDNA insert had a replication advantage in bacteria. Due to the increased diversity and functionality, we focused on the pMR766(+)L library.

The pMR766(+)L plasmid library showed deletions across the ZIKV genome (Fig. 3C), with a peak at NS1, and a region of zero deletion depth at the flanking region of the genome, corresponding to the plasmid backbone (ori/bla). Given that deletion variants from the pMR766(+)S, pMR766(–)L, and pMR766(–)S libraries did not ultimately passage efficiently in cells, we did not construct deletion landscapes for the libraries from these plasmids. Similar to the HIV plasmid landscape, deletion of these backbone regions compromises the ability of the plasmid to be propagated effectively in bacteria. The peak centered at NS1 reflected increased growth of mutants with deletion in NS1, which is known to have cryptic promoter activity and cause reduced growth in *E. coli* (60, 69, 70).

As with HIV-1, wild-type ZIKV and RanDeL variants were packaged by co-transfection of 293T cells using equal masses of each ZIKV RanDeL and the wild-type clone. Filtered, concentrated virus-containing supernatant was isolated, pooled, and used to infect Vero cells at high MOI (>16). The viral pool was passaged three times in Vero cells, in parallel to a wildtype-only control infection.

Viral RNA was analyzed from transfected 293T cells and cell-free supernatant at each passage by RT-qPCR to quantify RanDeL barcodes and total ZIKV genomes. First, we verified that barcoded mutants could be detected intracellularly post-transfection of each individual library (Fig S7A) and each day post-infection (dpi) in passage 1 (Fig. 3D). At 1 dpi, barcodes represented < 0.01% of total Zika genomes and did not increase in percentage by 3 dpi. Total viral genomes (ZIK-C) in RanDeL co-transfected samples were not significantly different from the control infection (Fig S7B), indicating the absence of a detectable interference effect from ZIKV variants, in agreement with the HIV results. However, a significant drop-off in barcode prevalence was observed between intracellular RNA post-transfection and supernatant RNA post-infection, indicating a strong selective pressure (i.e., bottleneck) on RanDeL variants between transcription and egress.

To identify CAEs, deletion-depth profiles were constructed by Illumina™ sequencing of ZIKV RanDeL barcodes after co-transfection, passage 1, and passage 2. The deletion landscape of intracellular RNA in co-transfected 293T cells was similar to the plasmid profile with a couple notable exceptions (Fig. 3E). First, at the 5’ end of the genome, deletions of the internal CMV promoter (bases 1–721) inhibited transcription because the CMV promoter is required for transcription of the ZIKV RNA genome. Second, at the 3’ end of the genome, deletions of the HDV ribozyme and poly(A) sequence also inhibited to transcription.

Next, we analyzed deletion landscapes from the serial passage in Vero cells. Although the pool of viral deletion mutants for infection was initially all four sub-libraries, less than 1% of detectable barcodes were from pMR766(-) libraries by the end of passage 2. The vast number of the observed barcodes (≈95%) were derived from the pMR766(+)L library, with the remaining (≈5%) from the pMR766(+)S library. One potential reason is that pMR766(-) genome replication was unable to be rescued by the wild-type virus supplying NS5 in trans.

Of the initial 40,000 mappable barcodes of pMR766(+)L, only 300 were detected after passage 1. These were used to construct a deletion profile (Fig. 3F), which shows a single peak, centered at *E*, that slopes downward in each direction to a deletion-depth of zero beyond the 5’ border of *PrM* and the 3’ border of *NS1*. Importantly, Pr, M, and E are 3 of the 4 structural proteins that comprise the viral particle (C is the last). By passage 2, several variants with 1-2 kb deletions that spanned this region increased 200–500× in prevalence. Flavivirus replicon systems have previously been developed by deletions in these regions, including C (41, 71). RanDeL-seq determined that ZIKV tolerated deletions in prM and E, but not in C.

A final deletion landscape of ZIKV by Passage 2 (Fig. 3G) showed that only 3 kb genomic interval of pMR766 can tolerate deletion. The region beginning exactly at Pr and ending precisely at the end of NS1 is TAE1, and can be efficiently complemented in *trans*. The regions flanking TAE1 are cis-acting, with CAE1 encompassing the 5’ UTR and CAE2 the remainder of the non-structural genes and 3’UTR (NS2-3’ UTR). Deletions within CAE1 or CAE2 were not detected upon passage. We verified these results by two serial passages in C6/36 cells (Fig S8).

## Discussion

We describe a high-throughput method (RanDeL-seq) to comprehensively map viral cis- and trans-elements at a single-nucleotide resolution. RanDeL-seq takes advantage of *in vitro* transposition, dual exonuclease chewback, and barcode cassettes to make randomly distributed deletions of varying size throughout a sequence of interest. As a proof-of-concept, we built and screened RanDeLs of >23,000 HIV-1 variants, and >90,000 ZIKV variants. Tracking and sequencing barcodes at each stage of the scheme (transfection and passage to passage) revealed elements critical for different stages in the viral life cycle, particularly transmission, and enabled mapping of these elements in HIV-1 and ZIKV at single-base resolution.

The deletion landscape for HIV-1 recapitulated known CAEs (LTR, Ψ, RRE) and their roles in transcription, encapsidation and egress. Despite previous claims that the Genomic RNA Packaging Enhancer (GRPE) is important for encapsidation (40), deletions of this region (nucleotide position 2022-2188) did not affect mobilization, in agreement with the findings of Nikolaitchik and Hu (72). The results also showed that the cPPT, despite its debated role in HIV-1 replication, was as important for sustained viral replication as the LTR, Ψ, and RRE. Surprisingly, we also identified a necessary 671-nucleotide region from the RRE to SA-7 which has not previously been reported and the function of which is undetermined. Previous work suggested that deletion to upstream cryptic splice sites 7a and 7b, but not SA-7, still allowed for HIV-1 replication (68). There were no instances of mutants where SA7 remained intact, and SA7a and 7b were deleted, as evidenced by the final HIV-1 deletion landscape. Additionally, SA-7 is heavily regulated, with an intronic splicing silencer (ISS), exonic splicing silencer (ESS), and exonic splicing enhancer (ESE). Interestingly, previous work suggested these elements were cis-acting (31, 73-75), but RanDeL-seq shows only the ISS included in CAE3, and therefore deletion of the ESS and ESE was tolerated or could be provided in *trans*.

Analysis of ZIKV deletion landscape, in two different cell types, showed that deletions in the C protein, non-structural genes NS2–NS5, and UTRs are not tolerated and could not be supplemented in *trans* by the wild-type virus. This ZIKV profile adds to established flavivirus CAE and TAE models that focus on conserved elements of 5’ and 3’ UTRs (76); as seen with Yellow Fever (41), West Nile Virus (77, 78), Dengue (79, 80), and Hepatitis C (81). While it accurately identifies the highly structured UTRs as critical CAEs, RanDeL-seq also demonstrated that deletions in C, NS5, and the other non-structural proteins (NS2-NS4) could not be complemented in *trans*. A recent study of the same strain of ZIKV reached similar conclusions through transposon insertion instead of deletion (82). Insertions in NS2-NS5 were not tolerated, except at the regions proximal to the protein cleavage sites. However, that group found that insertions were tolerated in the C protein, which RanDeL-seq labeled as a CAE.

We note a number of limitations to RanDeL-seq. First, the one-pot method can create extremely diverse libraries *in vitro*, but transformation of *E. coli* limits the library diversity, due to selection against potentially “toxic” or unstable sequences in viral genomes. This limitation is shown by the finding that the initial deletion depths are not flat across the viral genome. Specific regions (i.e. gag/pol HIV deletions) are still selected for despite having less coverage in the initial construction and transfection. RanDeL-seq may also be too inefficient to produce diverse libraries for viruses with much larger genomes, such as herpes viruses (encoded on 250 kb BACs) (83). Transformation of bacteria with high molecular weight DNA is inefficient, and large genomes are easily damaged by shearing during the physical manipulations required for cleanup. However, libraries could be developed by dividing large genomes into smaller pieces that can be mutated separately and then reassembled using suitable methods such as REXER (84) and could be incorporated into other elegant frameworks for mapping DIPs (85).

The HIV-1 screen was conducted using a single molecular clone of HIV-1 and a single clonal cell line (MT-4). It is possible that CAEs vary between viral strains and between cell lines and tissue types. Conducting the screen in a tissue explants (PBMC or HLAC cultures) may reveal different results. Also, the method is unable to monitor recombination between viruses (86), which could produce viral strains that have acquired more than one deletion, and create linkage effects. Similarly, no sequencing outside of the barcode cassette was done during serial passage, precluding the detection of additional mutations. However, we show a strong correlation between replicates, indicating that the observed selection was deterministic, rather than a result of drift.

Compared to pre-existing methods of CAE mapping, RanDeL-seq is able to cover the full viral genome with random deletions of variable size, track barcode (i.e. specific mutations) prevalence over time, and map at a single-nucleotide resolution. It is an improvement on methods of creating viral deletion mutants that rely on site-directed mutagenesis, iterative deletion, or spontaneous DI RNA emergence in culture; RanDeL-seq can comprehensively map full length viruses, not just one targeted location. Additionally, RanDeL-seq fully abrogates genomic regions, rather than silencing potential CAEs with SNPs, stop codons, or sequence changes that don’t affect protein synthesis. This full deletion allows determination of the essential nature of each genomic region.

The advantages of the method, along with its speed and low cost, make it attractive for studying novel, emerging viruses. The method can be rapidly deployed to identify CAEs for antiviral drug targeting, minimal sequences necessary for vaccine development, and candidates for novel antiviral therapies such as TIPs. Collectively, RanDeL-seq could be a valuable and versatile framework of general use to virology, aiding the study of viral replication mechanisms and the development of novel antiviral therapeutics.

## Materials and Methods

### Plasmids

pNL4-3 is a molecular clone of HIV-1 subtype B (87), and was a kind gift of Malcom Martin (AIDS Reagent Program #114). Two molecular clones of ZIKV, strain MR-766, were a generous gift from Matthew Evans. Two versions were available: a wild-type clone, pMR766(+), and a mutant, pMR766(-). The mutant clone has a GDD→GNN mutation in NS5 and lacks a functional RNA Dependent RNA Polymerase.

### Cells

All cells were grown at 37°C with 5% CO2. HEK 293T cells and C6/36 *Aedes albopictus* cells (American Type Culture Collection, # CRL-3216 and CRL-1660, respectively) were propagated in DMEM supplemented with 10% FBS (Fisher Scientific) and 1% Pen/Strep (Fisher Scientific), referred to as D10. Vero cells (African green monkey [Cercopithecus aethiops] kidney cells) (ATCC, # CCL-81) were also propagated in D10. MT-4 cells (NIH AIDS Reagent Program, #120) were propagated in RPMI 1640 supplemented with 10% FBS, 1% Pen/Strep, HEPES, and L-glutamine, referred to as R10.

### Reagent sourcing

All enzymes were obtained from New England Biolabs (NEB; Billercia, MA, USA) unless indicated otherwise. All chemicals were obtained from Sigma-Aldrich (St. Louis, MO, USA), unless indicated otherwise. DNA oligonucleotides and synthetic dsDNA were obtained from Integrated DNA Technologies (Coralville, IA, USA).

### Transposon DNA cassettes

Transposon cassettes were ordered in 3 pieces as synthetic dsDNA (<500 bp) (gBlocks, IDT) and cloned by Gibson Assembly into pUC19 (linearized at the BamHI site) (88). Post assembly, linear transposon cassettes were constructed by standard Q5 Hotstart PCR protocol (NEB) with TN5MK plasmid template and the oligos, oTN5-F and oTN5-R (IDT). The template was amplified under the following conditions: 98°C for 30 seconds; 15 cycles of: 98°C for 10 seconds, 68°C for 20 seconds, 72°C for 50 seconds; final extension: 72°C for 5 min, hold at 10° C. PCR products were purified by column with Zymo DCC-5 Kit (Zymo Research), then analyzed on a 0.8% agarose/TE gel. The 1.4 kb transposon cassettes were excised and cleaned using Qiagen Gel Extraction Kit (Qiagen) and Zymo DNA columns.

### Barcode DNA cassettes

HIV-1 barcodes were blunt-end, 5’-phosphorylated, 60 bp DNA cassettes prepared by standard Q5 Hotstart PCR with BC20v1-F and BC20v1-R on the oligonucleotide pool BC20-T (IDT). Oligos sequences can be sound in Table S2. BC20-T oligos were 60 bp ssDNA molecules with consensus 20 bp flanking sequences and a middle 20 bp with machine-mixed bases (random sequences for barcodes). Reactions were cycled at 98°C for 30 seconds; 5 cycles of: 98°C for 10 seconds, 65°C for 75 seconds; 1 cycle of: 65°C for 5 min, hold at 10° C. Post-PCR, barcode cassettes were column purified (Zymo). A 3’-dT overhang was added with a 3’→5’ exonuclease-deficient Klenow Fragment of *E. coli* DNA Polymerase I per manufacturer’s protocol. The reaction was incubated at 37°C for 3 hours. Post-incubation, DNA was cleaned by column-purification (Zymo) and eluted in Tris-acetate-EDTA buffer.

ZIKV libraries were prepared identically with a slight difference in the forward and reverse common sequences of the barcode cassette (BC20v2-F, BC20v2-R). These sequences modified a triple repeat in the forward barcode read (GGG) to avoid problems with sequencing of low-diversity libraries on the Illumina HiSeq4000.

### Chewback Conditions

Template DNA, λ-HindIII, was initially heated to 60°C for 3 minutes and immediately cooled to separate annealed cohesive cos ends. A standard 50 ul chewback reaction was prepared on ice by combining dH2O, 10X NEB2.1, λ-HindIII DNA template (500 ng/ul), T4 DNA Polymerase (3 U/ul), RecJf (30 U/ul), and ET SSB (500 ng/ul). The reaction was then incubated at 37° C. After 30 minutes, 1 ul of 10 mM dNTPs were added, the reaction mixed, and returned to 37°C for 11 minutes to allow T4 DNA Polymerase to fill in recessed ends. The reaction was halted by adding EDTA (pH 8.0) to a final concentration of 20 mM. Various dropout reactions were conducted, where dH20 was substituted for enzymes.

### Determination of Chewback Rate

A 4.3 kb dsDNA template was obtained by purifying the 4361 bp fragment of λ-HindIII digest. The λ-HindIII template was run out on a 0.8% agarose gel, stained with SYBR Safe, and excised. DNA was recovered by adding 0.1 gel volumes of β-agarase I reaction buffer (NEB), melting gel slices briefly at 65°C, cooling to 42°C, and immediately adding 1 U of β-agarase I per 100 ul of molten gel (NEB). The mixture was incubated at 42°C for 60 min to release DNA bound in the agarose matrix. DNA was precipitated from the digested fraction with sodium acetate (3M) and 2-propanol. After mixing, the reaction was spun at 20000×g for 15 min at 25°C, and the supernatant aspirated. The DNA pellet was washed once with 70% ethanol, allowed to air dry briefly, then dissolved in TE.

A chewback reaction was set up per minimal conditions and incubated at 37°C. At 0, 5, 10, 15, 20, 25, 30, 40, 50, 60, 70, and 80 minutes of incubation, an aliquot of the reaction was removed and combined with equal volume dNTP buffer (NEB2.1, 10mM dNTP, dH^2^ O). These 12 reactions were then incubated at 37°C for 11 min to allow T4 DNA Polymerase to fill in the single-stranded tails that remain uncleaved by RecJ^f^. After 11 min of fill-in, the reaction was halted with an equal volume of Stop Buffer (EDTA, dH2O).

The concentration of dsDNA was determined by a fluorimetric method (PicoGreen, Thermo Fisher Scientific). Each reaction was diluted in TE and mixed with a PicoGreen working stock (diluted to 1/200× in TE Buffer) to be read with an Enspire plate reader (Perkin Elmer) with 480 nm excitation and 520 nm emission filter. Fluorescence was compared to a λ DNA standard. All reactions were performed in triplicate. Chewback rates at 37°C were calculated by fitting the decay in dsDNA (fluorescence signal) at various timepoints to a linear regression model with the freely available R statistical software.

### Construction of RanDeL

#### DNA extraction, precipitation, and wash

Throughout construction of the random deletion libraries, DNA was extracted, precipitated and washed with the same methods. DNA samples were extracted with 25:24:1 phenol:chloroform:isoamyl alcohol equilibrated with TE, followed by a second extraction with pure chloroform (Sigma-Aldrich). The upper aqueous layer was transferred to a new DNA LoBind tube, and 25 ug of co-precipitating GenElute Linear Polyacrylamide (Sigma-Aldrich) added and the solution mixed to homogeneity. DNA samples were precipitated from the aqueous phase by MgCl2/PEG-8000 precipitation. Samples were adjusted to a final concentration of 12.5% (m/v) PEG-8000 and 20 mM MgCl2 by adding MgCl2 (1M) and 50% (m/v) PEG-8000. Reactions were inverted and flicked to mix, then spun at 20000×g for 60 minutes in a refrigerated microcentrifuge (Eppendorf) at 25°C to pellet all precipitated DNA. After centrifugation, supernatants were removed and discarded. Freshly prepared 70% ethanol was added and the reactions mixed by inversion. Samples were spun at 20000×g for 2 minutes to collect the pellet, then aspirate and discard the supernatant. Additional ethanol was added to wash the pellet, and samples were spun again at 20000×g for 2 minutes to collect DNA pellets. All supernatants were carefully removed and the pellet dried briefly at room temperature (5 minutes) until no visible liquid remained. DNA samples were solubilized by adding TE, incubating the tube at 42°C for 20 minutes and mixed by flicking the tube.

#### In Vitro Transposition

Transposon Cassettes were inserted into pNL4-3 by *in vitro* transposition with *EZ*-Tn5 transposase (Epicentre) per manufacturer’s protocol and with equal mols of plasmid template and transposon. After a two-hour incubation in reaction buffer at 37°C, the reaction was halted with 1% SDS solution and heated to 70°C for 10 minutes. The entire volume of the reaction was transferred onto a 0.025 um membrane floating on TE. Drop dialysis was allowed to proceed for 1 hour. Plasmids were electroporated into bacterial cells and selected with the encoded antibiotics (carbenicillin, kanamycin). Plasmid DNA was obtained by Qiagen Maxiprep according to manufacturer’s protocol.

#### Transposon Excision

Inserted transposons were excised by treatment with either meganuclease I-SceI or I-CeuI in CutSmart Buffer (NEB). Reactions were incubated at 37°C for 8 hours, with brief mixing by inversion performed every 2 h. DNA was extracted by phenol-chloroform, precipitated by MgCl2/PEG-8000, and ethanol washed for the next stages.

#### Chewback

Substrate DNA was heated to 60°C for 3 min and immediately placed on ice to separate DNA aggregates in preparation of chewback. Four standard chewback reactions were prepared, each with a different chewback length (5, 10, 15, and 20 minutes). At the appropriate time, the indicated reaction was removed from 37°C incubation and dNTPs were added. The reaction was mixed and returned to 37°C to allow T4 DNA Polymerase to fill in recessed ends. After 11 minutes of fill in, the reaction was halted and placed on ice. All chewback reactions were pooled and then extracted with two phenol chloroform extractions. The DNA was desalted by running through separate Sephacryl gel filtration columns (Microspin S-400 HR columns (GE Lifesciences)).

#### End Repair

DNA was pooled and blunt-ended by NEBNext End Repair Reaction Module (NEB), incubating at 20°C for 30 minutes. DNA was extracted by phenol-chloroform, precipitated by MgCl2/PEG-8000, and ethanol washed for the next stages.

#### Addition of 3’-dA overhang to backbone

A 3’-dA overhang was added to the purified blunt-end truncated linear pNL4-3 DNA with a 3’→5’ exonuclease-deficient Klenow Fragment of E. coli DNA Polymerase I (NEB). The reaction was incubated at 37°C for 1 hour, and then heat-inactivated (70°C for 20 minutes). Treatment with Antarctic Phosphatase (NEB) per manufacturer’s protocol dephosphorylated the 5’ ends (1 hour at 37°C, 5 minutes at 70°C to deactivate). DNA was migrated on a 0.8% agarose gel and stained with SYBR Safe. All DNA vectors greater than 8 kb were excised, recovered with β-agarase, precipitated with sodium acetate and 2-propanol, and ethanol washed.

#### Ligation of Barcode Cassettes and chewed vector

3’-dT-tailed barcode cassettes were ligated into a 3’-dA-tailed vector and the DNA circularized using T4 DNA Ligase in a PEG-6000 containing buffer (Quick Ligation Buffer, NEB). Ligation was performed at a 30:1 insert:vector molar ratio at bench temperature (24°C) for 2 hours, then the reaction was halted by adding EDTA (pH 8.0) and mixing. Next, Proteinase K (800 U/ml) (NEB) was added, the reaction mixed, then incubated for 30 min at 37°C to cleave bound T4 DNA Ligase from the DNA.

#### Sealing of nicks in hemiligated DNA

Nicked DNA was sealed by sequential treatment with T4 Polynucleotide Kinase (T4 PNK) and *Taq* DNA Ligase. Hemiligated DNA was 5’-phosphorylated with T4 PNK in T4 DNA Ligase Reaction Buffer (NEB) at 25°C for 30 minutes. Reactions were purified with AMPure XP beads (Beckman-Coulter Genomics) and eluted with T4 DNA Ligase Master Mix. The eluate was incubated at 37°C for 60 minutes to phosphorylate DNA at the nicked sites. The nicks were then sealed by treatment with Taq DNA Ligase (NEB) in Taq DNA Ligase Reaction Buffer at 75°C for 15 minutes. Ligated DNA was purified by AMPure XP beads and eluted in TE.

#### Library transformation and outgrowth

The purified ligation was electroporated into electrocompetent *E. coli* (DH10B) cells. Cells were allowed to recover in SOC (Thermo Fisher), and then expanded for overnight growth in LB-Miller supplemented with carbenicillin. Finally, Deletion Library Plasmid DNA was isolated from spun-down, harvested cultures by Qiagen Maxiprep.

### Transfection of Viral Stocks

Co-transfections were with an equal ratio of wild-type and deletion library plasmid. 293T cells were added to flasks at a ratio of 5e6 cells/mL in DMEM supplemented with 25mM HEPES. Wild-type and deletion library plasmids were diluted in unsupplemented DMEM (i.e., no serum or antibiotics added) to a concentration of 10 ng/mL total DNA and PEI was added to a concentration of 30 µg/mL in a volume ∼10% of the total volume in the transfection well or dish (e.g., 200 µl for a 6-well plate with 2 mL media). The transfection mix was vortexed, incubated at bench temperature (24°C) for 15 minutes, then added to the 293T flasks. Media was replaced after an overnight incubation (16–20 hours). Virus was harvested at either 48 hours (HIV) or 72 hours (ZIKV) post-transfection by passing through 0.45 μm sterile filters (Millipore). HIV-1 stocks were prepared with pNL4-3 and the pNL4-3 deletion library. ZIKV viral stocks were prepared with pMR766(+) and one of the four MR-766 deletion libraries: pMR766(+)ΔS, pMR766(+)ΔL, pMR766(-)ΔS, or pMR766(-)ΔL.

### HIV High MOI Screen

#### Concentration of Virus

Concentrated virus was prepared by ultracentrifugation (Beckman Coulter Optima XE-90, rotor SW 28) at 20,000 rpm through a 6% iodixanol gradient (Sigma Aldrich, D1556-250mL) for 1.5-2 hours at 4°C.

#### Titration of Viral Stocks

The concentrated HIV-1 stocks were titrated by infecting cultures of MT-4 with concentrated virus and scoring for HIV p24-producing cells at 24 hours post-infection. Virus was added to MT-4 cells in R10, mixed briefly, then incubated for 4 hours at 37°C. After 4 hours, additional media was added, and the infection allowed to proceed for an additional 20 hours (a single-round of replication). At 24 hours post-infection, cultures were fixed with 20% formaldehyde (tousimis) and incubated for at least 1 hour at 4°C. After fixing, cells were permeabilized by treatment with 75% ice-cold methanol for 10 minutes, then stained with a phycoerythrin-labelled monoclonal antibody (KC57-RD1,BD) for 30 minutes before washing once in stain buffer. At least 50,000 live cells were counted by flow cytometry on a FACS Calibur DxP8. Gates were drawn based upon stained naive cell population. Analysis was done in FlowJo.

#### High MOI passage scheme

On day 0, 2*10^6^ MT-4 cells were infected at a MOI of 5-20 with the prepared and titrated virus pool for 4 hours in a volume of 2 ml, then transferred to a T25 flask containing 10 ml of MT-4 cells at a concentration of 10^6^ cells/ml. On day 2 (40 hours post infection (hpi)), the 12 ml of culture was transferred to a T175 flask containing 60 ml of MT-4 cells in R10 at a concentration of 10^6^cells/ml. On day 3 (70-72 hpi), supernatant from the MT-4 was clarified by centrifugation and 0.45 μm filtration, then concentrated by ultracentrifugation as described above. One cycle corresponds to 3 rounds of HIV-1 replication (completed on day 1, day 2, day 3) and was repeated four times for a total of 12 passages (i.e. rounds of replication). The cycle was repeated a total of four times (12 passages / rounds of replication) with 3 biological replicates (K, L, M). Wild-type pNL4-3 controls were passaged alongside the deletion library, also in triplicate (A, B, C).

#### Viral RNA Isolation

Viral RNA was isolated from the concentrated virus pool at passage 0, passage 3, passage 6, passage 9, and passage 12 using a QIAmp Viral RNA Mini Kit (Qiagen) per the manufacturer’s instructions with two exceptions: 1) carrier RNA was replaced with 5 of linear polyacrylamide (Sigma) per isolation; and 2) 5·10^6 copies of bacteriophage MS2 RNA (Roche) were spiked in per isolation. Total cellular RNA from 293T cells was isolated using Trizol (Life Technologies) from cell pellets obtained at the time of viral harvest. A poly(A) fraction, representing mRNA, was isolated by annealing total RNA to magnetic d(T)25 beads to pull down polyadenylated transcripts (NEBNext Poly(A) mRNA magnetic isolation module).

#### RT-qPCR Analysis

Purified vRNA was reverse-transcribed with Superscript III (Thermo Fisher) and Random Primer Mix (New England Biolabs) for quantification by RT-qPCR with Fast SYBR Green Master Mix (Thermo Fisher). Barcode cassettes were quantified by oligos BC20v1-F and BC20v1-R. Total HIV RNA was estimated by primers targeting HIV *pol*, NL43pol-F and NL43pol-R. Samples were normalized for recovery by determining levels of MS2 RNA recovered by oligos MS2-F and MS2-R (sequences from (89)). Relative expression was calculated by traditional RT-qPCR methods (90). Oligos sequences can be sound in Table S2.

### ZIKV High MOI Screen

#### Concentration of Viral stocks

Virus stocks were concentrated by ultrafiltration. Clarified supernatant was added to a 100 kDa MWCO filtration device in 20 aliquots. The device was spun at 1200×g for 20–30 min until the concentrate volume was less than 1 mL. The flowthrough fraction was removed and an additional supernatant added to the upper reservoir and the process repeated. Generally, clarified supernatant was concentrated 20–40X. Concentrated stocks were adjusted to 20% (v/v) FBS and 10 mM HEPES (to reduce loss in infectivity from freeze-thawing).

#### Titration of Viral Stocks

ZIKV stocks were titrated by plaque assay (91). On the day before infection, Vero cells were seeded in 6-well or 12-plates at approximate 50% confluency. On the day of infection (0 dpi), serial 10-fold dilutions of sample stocks were prepared by dilution in DMEM supplemented with 3% (v/v) heat-inactivated FBS. The media from each well of the infection plate was removed and replaced with serially-diluted virus. The plate was gently rocked and returned to the incubator for a period of 1 hour, with gently rocking applied every 15 minutes. After one hour of adsorption, the virus was removed and the cultures overlaid with a viscous solution of 1% (w/v) carboxymethylcellulose (Sigma #C4888) in DMEM-F12 (8% FBS, 1% pen/strep). Infection plates were returned to the incubator and left undisturbed for 5 days. At 5 dpi, the wells were with 20% formaldehyde and mixed gently for 1 hour. The supernatant was removed and the culture stained with a solution of 1% crystal violet in 20% ethanol for 15 minutes. Wells were de-stained by rinsing with dH2O. Plaques were 1-2 mm in diameter and could be visualized as clear circular patches on the stained purple monolayer.

#### High MOI passage scheme

On day 0, Vero cells were infected at a MOI of 16–30 with a virus pool containing wild-type ZIKV and ZIKV deletion libraries. The inoculum was applied in a low volume in a 6-well plate for 1 hour, then removed. Supernatant was collected at 1, 2, and 3 dpi, corresponding to one passage. Virus from passage 1 was titrated by plaque assay and used to infect Vero cells for passage 2. The passage scheme was conducted with 2 biological replicates.

#### Viral RNA Isolation

ZIKV Viral RNA was isolated from the concentrated virus pool at 293T transfection, passage 1, and passage 2 per similar methods to the HIV screen.

#### RT-qPCR Analysis

Purified RNA was reverse-transcribed with MuLV-R (NEB) and Random Primer Mix (NEB) for quantification by RT-qPCR with SYBR Green Master Mix. Barcode cassettes were quantified by oligos BC20v2-F and BC20v2-R. Total ZIKV RNA was estimated by primers targeting the ZIKV capsid protein (ZIK-C), MR766-C-F and MR766-C-R. Samples were normalized for recovery by determining levels of MS2 RNA recovered by oligos MS2-F and MS2-R. Relative expression was calculated as done in the HIV-1 screen. Oligos sequences can be found in Table S2.

### NGS Analysis

#### Genotyping of plasmid libraries

Insertion and deletion plasmid libraries were prepared for paired-end sequencing on the Illumina HiSeq/MiSeq platforms by a Nextera XT Kit (Illumina) from 1 ng of each library. Transposon insertion and PCR enrichment were performed per the manufacturer’s instructions, but the sublibraries were pooled and size-selected by running out on a 1.5% agarose gel, staining with SYBR Safe (Thermo Fisher), and excising a gel fragment corresponding to DNA of size range of 350–500 bp. DNA was purified from the gel slice using Qiagen Buffer QG, Buffer PE (Qiagen), and DCC-5 columns (Zymo Research). The sublibraries were pooled and sequenced on a single lane of a HiSeq4000 (Illumina), using 2×125 bp reads.

Transposon insertion locations were computed by filtering for high-quality reads containing an exact match of either mosaic end sequence of TN5MK, then extracting flanking regions to build an insertion map. A lookup table matching deletion locus to barcode sequence was determined by: 1) searching reads for the forward and reverse common barcode sequences and extracting the intermediate 20 bp; 2) assembling a list of barcode sequences; and 3) assigning flanking regions to each barcoded deletion using custom Python software.

#### Sequencing of serial passage

Illumina sequencing libraries were prepared by a modification of a method specified in (92) and detailed in Figure S5. Barcode cassettes were amplified using a minimum number of cycles (typically 12-18) to prevent overamplification (post log-phase PCR). Illumina adaptors were added by two rounds of PCR (5 cycles each), to add phasing adaptors, random barcodes, and multiplexing barcodes. Sublibraries were size-selected on 5% TBE polyacrylamide gels and pooled for sequencing. 20–30 sublibraries were sequenced on two lanes of a HiSeq4000 (Illumina) (spiked with 25% PhiX), using a single 1×50 b read at the Center for Advanced Technology at University of California, San Francisco. Barcodes were tallied using custom Python software and matched to deletion loci using the lookup table prepared previously to calculate deletion depth.

## Data availability

The data that support the findings of this study are available from the corresponding author upon reasonable request.

## Biological materials availability

All unique biological materials are available from the corresponding author.

## Code availability

Custom code is available upon request.

## Acknowledgements

We thank the Weinberger lab for discussions and suggestions. We thank Kathryn Claiborn for editing. The following reagents were obtained through the NIH AIDS Reagent Program, Division of AIDS, NIAID, NIH: MT-4 from Dr. Douglas Richman (cat# 120) and pNL4-3 molecular clone from Dr. Malcolm Martin (cat #114). This work was supported by the Bowes Distinguished Professorship, the Alfred P. Sloan Research Fellowship, the Pew Scholars in the Biomedical Sciences Program, the DARPA INTERCEPT program (D17AC00009), and the NIH Director’s New Innovator (OD006677) and Pioneer Award (OD17181) programs.

## Author contributions

T.N. and L.S.W. conceived and designed the study. T.N., V.R.S, and C.E.T., designed and performed the experiments, and curated the data. J.J.G. and L.S.W. wrote the paper.

## Competing interests

The authors declare that they have no competing interests

Supplementary information is included with this manuscript

